# Controlling cluster size in 2D phase-separating binary mixtures with specific interactions

**DOI:** 10.1101/2022.02.10.479877

**Authors:** Ivan Palaia, Anđela Šarić

## Abstract

By varying the concentration of molecules in the cytoplasm or on the membrane, cells can induce the formation of condensates and liquid droplets through phase separation. Their thermodynamics, much studied, depends on the mutual interactions between microscopic constituents. Here, we focus on the kinetics and size control of 2D clusters, forming on membranes. Using molecular dynamics of patchy colloids, we model a system of two species of proteins, giving origin to specific heterotypic bonds. We find that concentrations, together with valence and bond strength, control both the size and the growth time rate of the clusters. In particular, if one species is in large excess, it gradually saturates the binding sites of the other species: the system becomes then kinetically arrested and cluster coarsening slows down or stops, thus yielding effective size selection. This phenomenology is observed both in solid and fluid clusters, which feature additional generic homotypic interactions and are reminiscent of the ones observed on biological membranes.

## I. INTRODUCTION

Phase separation is a widely present phenomenon in cells^1,2^. When concentrations of certain biochemical species exceed a threshold, condensates start to form, which bring together one or more macromolecules present in the cytoplasm. The stability of these condensates relies on the mutual interactions between the species involved: interactions can be either generic, as for proteins’ intrinsically disordered tails coming together because of hydrophobicity, or specific, as for amino acid groups that form physico-chemical bonds only with given other groups^3–5^. In addition, attractive interaction or binding can occur between two specimens of the same species (homotypic interaction) or between different species (heterotypic interaction). Much progress has been made in understanding the phase diagrams of many of these systems, which determine whether the mixed or the demixed phases are stable at equilibrium, both from the experimental and the theoretical point of view^6,7^.

Condensates, or clusters, of biomolecules not only form in the cytoplasm, but also on two-dimensional membrane surfaces^8–12^. This occurs when at least one of the species involved is embedded in the membrane: for instance, the clustering of transmembrane protein LAT activates signalling to produce cytokines when T-cell receptors are exposed to an antigen^9^, while protein Whi3 in fungi condenses with RNA on the endoplasmic reticulum^10^.

Increasing density is often assumed to promote the creation of a dense phase, thus favoring phase separation. However, re-entrant behaviour of the dense phase with respect to concentration has been observed in several phase-separating systems, suggesting that increasing concentration of a given species does not necessarily promote the formation of large assemblies. While in some cases there appears to be an electrostatic-related change in the interactions^13–15^, in others the effect seems to be fully stoichiometric^16–19^ and most likely functional^20^.

Computational work on biological condensates has built upon various models, ranging from simple lattice models^21,22^, to off-lattice coarse-grained polymers^23,24^, stickers and spacers^25,26^, or patchy colloids^17–19,22,27^. In many studies, though, self-assembly is represented within the scaffold-client context, which assumes the existence of a set of proteins (scaffold) that can alone phase-separate through homotypic interactions^2,19,27,28^. While many systems correspond to this descriptions, others, especially in 2D, do not and feature mainly specific heterotypic interactions^3,5,17^.

While a large effort has been directed toward the understanding of the thermodynamics of phase-separating systems, the kinetics of aggregation, namely the mechanisms of growth and coarsening of clusters, has been studied less. Predictions often rely on classical kinetic theory^29–34^, oblivious of biological systems’ specificity, and whose validity needs to be tested on a case-by-case basis. In a dynamic environment such as that of a cell, where thermodynamic equilibrium is rarely reached, a better understanding of aggregation kinetics could provide new tools to extract information on the microscopic components from macroscopic experimental observations^10,35–37^.

In this paper, we study the kinetics of aggregation of a binary mixture with heterotypic interactions. The two species, that we name A and B, represent two kinds of proteins that can freely diffuse in a 2D environment, such as a biological membrane. We describe such proteins as patchy colloids, endowed with binding sites that can form bonds with binding sites of the opposite species (Fig. 1): an A and a B particle can form a bond, but two A or two B particles cannot. By geometrical constraints, bonds are exclusive, meaning that a bond cannot connect more than two binding sites and a binding site already participating in a bond cannot engage in a second bond. The number of binding sites of either species is the valence, that we denote by *q*_A_ and *q*_B_. For simplicity we restrict ourselves to the equal-valences case *q*_A_ = *q*_B_ = *q*. We study a range of integer valences *q* (between 3 and 5), a range of concentration ratios between

**FIG. 1.**
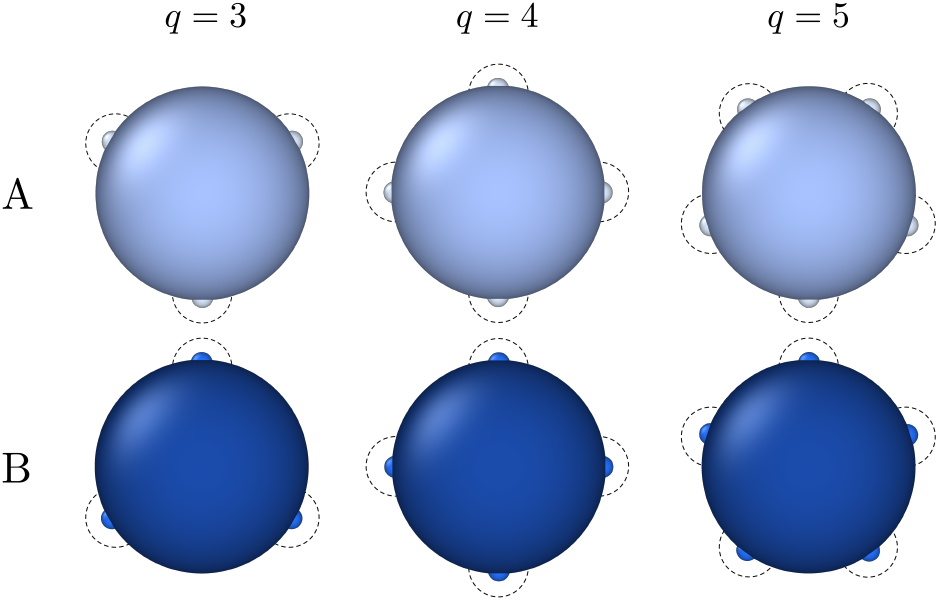
Sketch of particles of type A (violet) and type B (blue) for valence *q* = 3, 4, and 5. The big violet/blue disks, of radius *σ*, represent volume exclusion. Patches on A particles interact only with patches on B particles: when two interacting patches, of radius 0.1*σ*, superimpose, the energy gain is maximum and equals −*ε*. The dashed lines delimit the interaction area, of radius 0.3 *σ* : two patches attract each other only if their interaction areas intersect. See Methods for details.

B and A particles, and a range of different bond energies. For each system, at given values of valences, concentrations and bond energy, we perform Molecular Dynamics (MD) simulations and observe the self-assembly of clusters of A and B particles, of various composition and size. We find that concentration, valence and bond strength influence the rate of coarsening of the cluster, and in particular can drive the system to a kinetically arrested state, effectively resulting in cluster size selection.

## II. METHODS

In our model of patchy particles, volume exclusion between particles is enforced through the Weeks-Chandler-Anderson potential

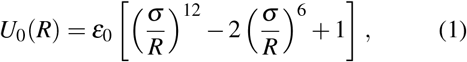

where *R* is the distance between the centres of the two particles, *σ* is their diameter, and *ε*_0_ is chosen equal to 10 *kT*. Patches are represented by ghost atoms that move rigidly with the particle; they are positioned at a fixed angular distance of 360°*/q* from each other (as in Fig. 1) and at a radial distance 0.475 *σ* from the centre of the particle. Patches on an A molecule interact only with patches on a B molecule, with the following attractive potential:

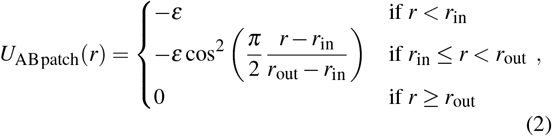

where *r* is the distance between the centres of the two patches, *ε* is the bond strength and is a parameter that we vary, *r*_in_ = 0.05 *σ* is the range within which interaction energy is strongest, and *r*_out_ = 0.15 *σ* is the range of attraction. We consider that two particles of type A and B form a bond when they have two patches at a distance *r < r*_out_. The chosen geometry and parameters finely prevent an A molecule from forming more than one bond with the same B molecule, and viceversa.

Simulations are run in LAMMPS^38,39^ and visualised with OVITO^40^. We simulate a number of A particles *n*_A_ equal to 200 and vary the number of B particles *n*_B_ to produce the desired concentration ratio. At the beginning of each simulation, particles are positioned at random sites of a square lattice, defined such that the density of A particles is always equal to 0.03 *σ* ^−2^. This value of density is physiologically plausible for transmembrane proteins, which often bind to cytoplasmic proteins of variable density, localised close to the membrane^17^. The timestep chosen for the integration of the 2D equations of motion is *τ*_*s*_ = 0.01 *τ*_0_, where 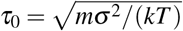 is the simulation unit of time and *m* is the mass of an atom (a whole patchy particle has mass (*q* + 1)*m*). Periodic boundary conditions are applied along both Cartesian directions. Every simulations is run for 5 × 10^7^ time steps *τ*_*s*_, a time large enough to allow cluster growth and nucleation, when present. To mimick an implicit solvent, the system is subjected to a Langevin thermostat of friction coefficient *γ* = 1 *m/τ*_0_. Finally, every point in the plots shown is an average over at least 20 realisations of the simulation, run with different random number generator seeds.

We analyse selected time steps with a clustering method based on the computation of an adjacency matrix. The average cluster size (Figs. 3, 6 and 8) is defined as the number of A molecules present per cluster, averaged over clusters containing at least 2 A and 2 B molecules, from several realisations of the simulation. The fraction of single-bound clustered particles (Fig. 4) is defined as the proportion of single-bound particles versus total number of particles in a given cluster, averaged over all clusters bigger than 30 particles, from different realisations. For all these variables, error bars represent standard deviation of the averages.

We quantify compactness of clusters (plots not shown), by computing the moment of inertia of each cluster relative to an in-plane rotation about its center of mass; we then normalise such value by the analogous moment of inertia for an elliptic cluster of similar aspect ratio and of uniform density, equal to that of fully packed hard circles. The inverse of the obtained quantity defines our compactness. This procedure was already used and thoroughly described in Ref. 17.

Clusters are made fluid (this refers only to Fig. 8) by adding to the repulsive potential (1) an isotropic attraction between A-A, A-B, and B-B particles, with the following shape:

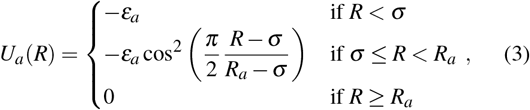

where *R* is again the distance between the centre of the two particles, *ε*_*a*_ is the isotropic attraction strength, and *R*_*a*_ is the isotropic attraction range.

## III. RESULTS

### A. Valence, bond energy and molar ratio control cluster size

During the simulations, we observe the formation of clusters of A and B particles, some of which are represented at the bottom of Fig. 3. Clusters look roughly crystalline for *q* = 3 (triangular lattice) and *q* = 4 (square lattice), while they have an amorphous structure for *q* = 5, in which case volume exclusion forbids that all 5 bonds be satisfied. This gives us the opportunity to study the effect of valence both in crystalline condensates, which are of simpler understanding and are akin to bidimensional pathological aggregates^41^, and in amorphous condensates, more likely relevant for functional phase-separation phenomena occurring on the membrane.

We are interested in the effect of stoichiometry on phase separation, therefore we fix the number of A particles *n*_A_ in our simulation box (whose surface is constant) and vary the number of B particles *n*_B_ from 0.25 *n*_A_ to 4 *n*_A_.

The average connectivity of an A particle, defined as the number of bonds formed on average by an A particle and ranging from 0 to *q*, is represented in Fig. 2 for the last time frame of our simulations. For all valences, connectivity increases as we increase the relative concentration of B particles *n*_B_*/n*_A_. Indeed, A molecules cannot condensate alone and need B molecules to act as crosslinkers between them.

**FIG. 2.**
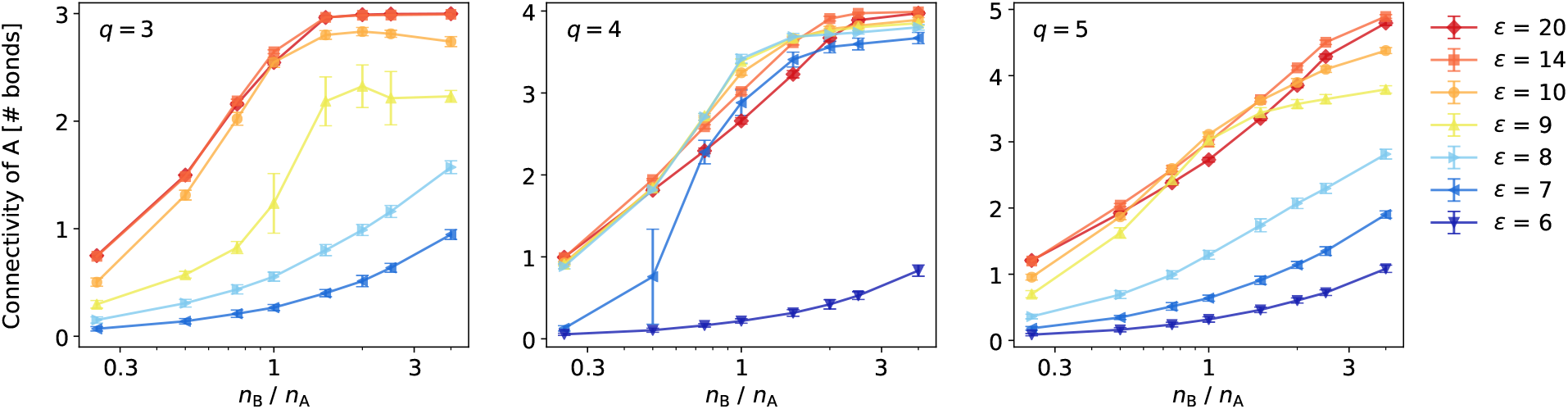
Average connectivity of A molecules, at different binding energy values *ε* (in units of *kT*), for *q* = 3, 4 and 5, at the end of the simulation.

To better quantify the extent of clustering, which would be completely but unpractically described by the full cluster size distribution, we compute the mean of such distribution, for the last time frame of our simulations (see Methods). The effect of the concentration of B linkers on the average cluster size is represented in Fig. 3, for different values of valence *q* and bond energy *ε*. Firstly, the plots show that clusters do not form within simulation time (or do not grow) if the binding energy is below a certain threshold, depending on *q*. Such threshold is approximately 8 and 6 *kT* for *q* = 3 and 4, respectively. This is because in the dense phase bonds have to provide a certain energy to win the entropy of the dilute phase: in the bulk, *q* bonds contribute to such energy, so the necessary energy per bond is inversely proportional to *q*. For *q* = 5, the threshold bond strength increases back to 8 *kT*, because the amorphous dense phase is frustrated and exhibits on average less than 4 bonds per molecule. As in 3D, phase separation is therefore naturally enhanced by valence (provided that full valence is physically achievable) or at least by the effective valence exhibited in the dense phase^27^.

Most interestingly, Fig. 3 shows that clustering is non-monotonic in the concentration of cross-linkers B. This is true at any *q* and at any *ε* above clustering threshold. Indeed, if there are too few cross-linkers, it is impossible to link all A molecules, some of which will remain free. Increasing the molar ratio *n*_B_*/n*_A_, up to 1, thus promotes both A connectivity (Fig. 2) and cluster size (Fig. 3). However, if cross-linkers are overabundant, they compete for binding sites on A molecules and end up filling all of them. It becomes then impossible for an already bound B molecule to find an empty binding site from another A molecule, so that A molecules become coated by B molecules and cross-links do not form (see cluster snapshots in Fig. 3). Average cluster size decreases, while A connectivity tends to saturate to *q*. A signature of this phenomenon is the average fraction of capping particles within a cluster, i.e. the proportion of clustered particles that are involved only in a single bond: this number is minimum at *n*_B_*/n*_A_ = 1, and increases as the molar ratio deviates from 1 (Fig. 4).

**FIG. 3.**
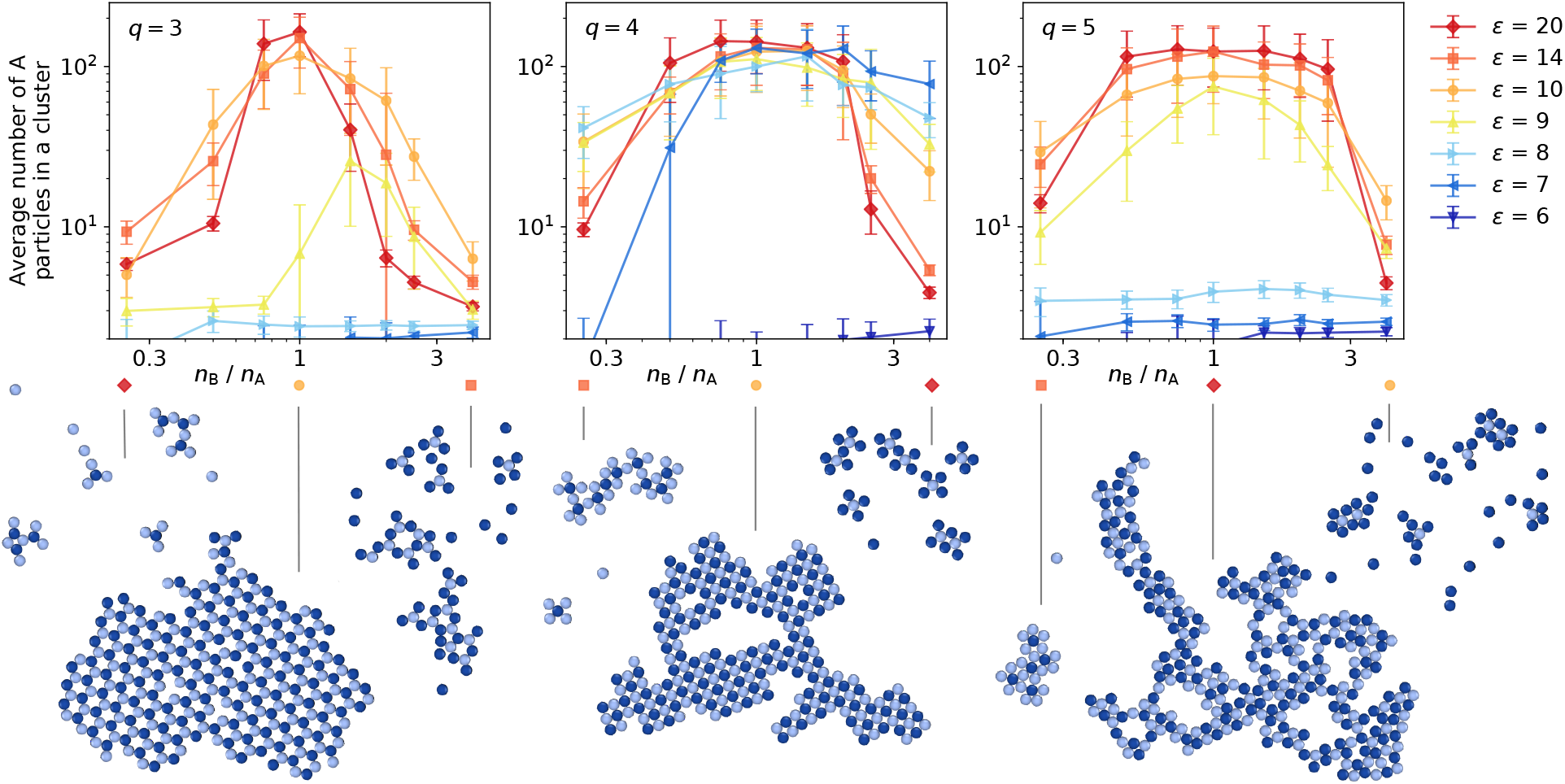
Average cluster size, at different binding energy values *ε* (in units of *kT*), for *q* = 3, 4 and 5, at the end of the simulation. Only clusters bigger than 4 molecules are counted (see Methods). Below, for each valence, snapshots of typical clusters are shown for concentration ratios of 1/4, 1 and 4 (pointed at by vertical lines), and for chosen *ε* (indicated by the symbols).

**FIG. 4.**
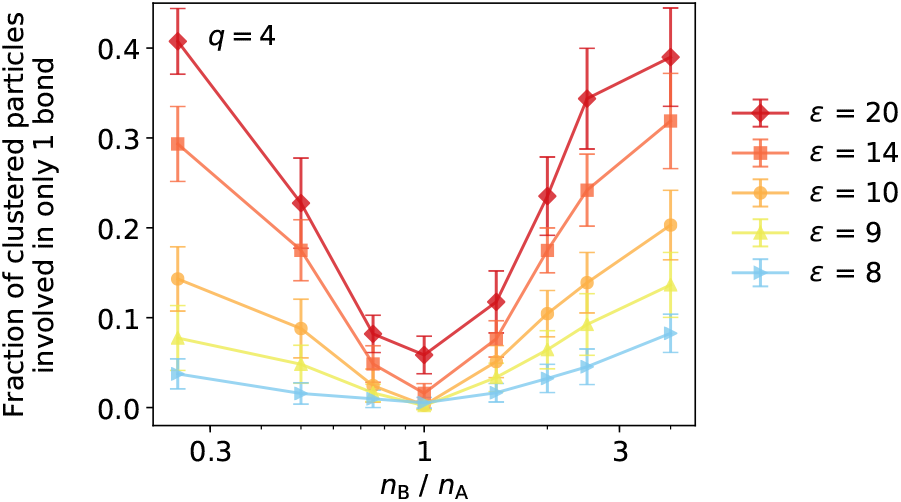
Fraction of clustered particles involved in one bond only, at the end of the simulation, for *q* = 4, at different binding energy values *ε* (in units of *kT*). Valences *q* = 3 and 5 exhibit similar behaviour.

In Figs. 3 and 4, the non-monotonic behaviour is more pronounced at strong than at weak binding *ε*. This is because at strong binding, bonds are irreversible and as soon as a full B cap forms around two given A molecules, no link will ever form between them. Instead, when binding is weak, caps have a finite lifetime and B molecules can detach, thus allowing, to some extent, for bond reorganisation and formation of new links; this favours the formation of larger clusters and of a proper dense phase. In other words, the cluster size is non-monotonic not only in the molar ratio *n*_B_*/n*_A_, but also in the bond energy *ε* (at least far from equal molar ratio): if *ε* is too low, clusters do not form because the energy gain is not sufficient to overcome the higher entropy of the dilute phase, but if *ε* is too large, the system gets stuck in a state where capping prevents coarsening.

Particular attention needs to be devoted to the equal concentration case, where we observe a non-monotonicity in cluster size with *ε* as well. Such phenomenon is analogous to magic-number effects reported in previous studies of many 3D systems or models^21–24^, where fine tuning of valence and/or concentration can cause major variations in the condensate properties. This is seen in Fig. 2, particularly in the *q* = 4 and 5 cases, where connectivity is enhanced at *n*_B_*/n*_A_ = 1 for inter-mediate bond strengths *ε*, but hampered for low or strong *ε*. This is again because lower *ε* allows for more bond reorganisation, indeed clusters at intermediate *ε* appear more compact and with fewer defects (see clusters snapshots in Fig. 2 at different bond strengths, and the Methods for a quantification of compactness). This is particularly important at equal molar ratio, when, due to the limited amount of B, full A connectivity can only be reached if no defects are present in the dense phase and the cluster is as compact as possible. At *n*_B_*/n*_A_ ≫1, this phenomenon is not relevant anymore, because the linker B is overabundant and A sites can be saturated anyway, by forming small B-coated clusters as described above; in such case, lower *ε* just means lower bond life time, and thus lower A connectivity (as shown by the right-handside parts of the panels in Fig. 2).

### B. Arrested state or thermodynamic equilibrium?

On can wonder whether the fully saturated state we observe at large concentration ratios is due to kinetic arrest or is a thermodynamic minimum. To gain physical intuition, we compare the free energy of two limit states, sketched in Fig. 5. For simplicity we discretise space and imagine *V* sites, filled by a number *n*_A_ of A particles and a number *n*_B_ *> n*_A_ of B particles; the remaining sites are left empty.

**FIG. 5.**
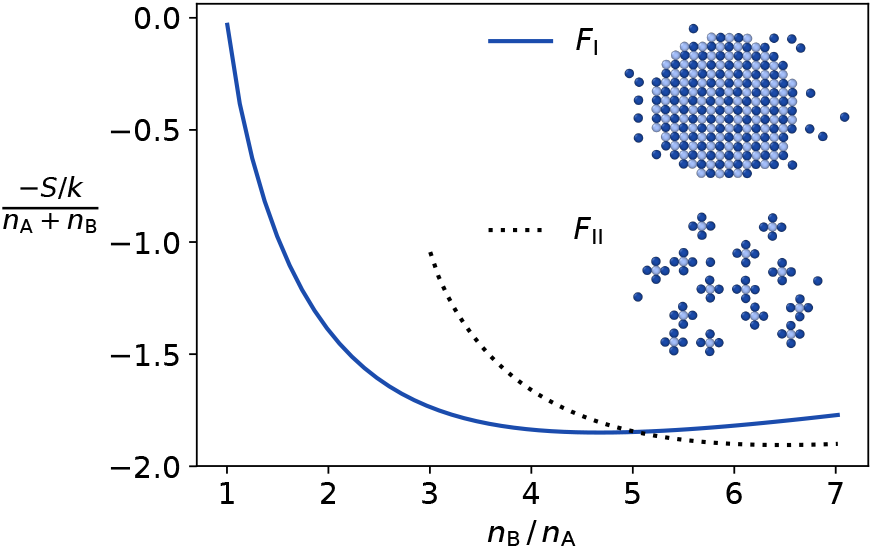
Entropy *S* of the lattice model, at *q* = 4, for the two limit cases: I, where all A particles form a single cluster, and II, where each A particle forms an independent cluster with 4 B particles.

A limit state (state I) is the one featuring one maximally connected, giant, round cluster with *n*_A_ A particles and ∼*n*_A_ B particles, while ∼ *n*_B_ − *n*_A_ B particles are free (i.e. they form the dilute phase). Its free energy, for low densities and strong bonds, can be approximated as

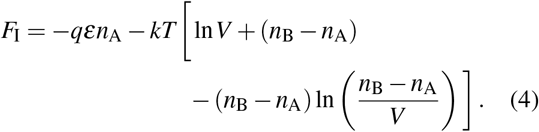

The first term in square brackets refers to the translational entropy of the cluster, while the last two terms represent the translational entropy of the free B particles.

The second limit state (state II) is the one where all A particles are fully capped by otherwise unbound B particles, so that the system has *n*_A_ tiny clusters, each made of 1 A and *q* B particles, and *n*_B_− *qn*_A_ free B particles. Clearly, this scenario is only possible if *n*_B_ > *qn*_A_. The related free energy is approximately

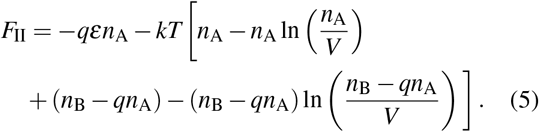

The first two terms in square brackets represent the entropy of the tiny clusters, while the last two the entropy of the free B particles.

Between states I and II, the one with the lower free energy is the thermodynamically stable one. Note that both states have the same energy (first term in both equations) as all A particles have their binding sites filled, so that entropy tips the scale. Entropy for states I and II is represented in Fig. 5. The single cluster (solid line) is still substantially favoured at *n*_B_*/n*_A_ = *q*. At larger *n*_B_, the entropic advantage decreases and then reverses. However the free energies of the two states remain comparable on the scale of a *kT* per molecule.

In our definition of state I, we considered the giant cluster to have a defined (circular) shape. While making the calculation easier, this eliminates from the count many configurations of equal energy − *qεn*_A_ and featuring one single cluster. If we redefine state I to account for all possible shapes, we can instead interpret *F*_I_ as a constrained free energy limited to minimal-energy configurations with one single cluster (just like *F*_II_ is a constrained free energy limited to configurations with *n*_A_ clusters). This leads to a new expression for *F*_I_, worked out in the Appendix. In short, non-circular shapes require more B particles to coat the perimeter of the cluster: while enlarging the configuration space for the giant cluster, this reduces the number of free B particles and therefore the associated entropy. Using results from the statistics of lattice animals^42–45^, we can group configurations by cluster perimeter and find that the cluster of minimal perimeter is the most favourable one, so that Eq. (4) is still approximately valid for our redefined state I.

Altogether, this suggests that the fully-capped state (II), while prevalent in our simulations, is not necessarily thermodynamically stable and that some kinetic lock is essential to make such a state the observed one at large concentration ratios.

### C. Kinetics of clustering

Prompted by the above considerations, we analyze the kinetics of clustering, which are likely relevant precisely for the size-control mechanism described in section III A. The time evolution of the average cluster size is represented for *q* = 4 in Fig. 6, for three different molar ratios. At equal concentration (middle panel), the number of particles in the clusters (i.e. their surface) steadily grows with time for all interaction energies *ε*, provided that enough time is allowed for nucleation at intermediate *ε*. A nucleation process is particularly evident for *ε* = 7 *kT*, where the curve stays flat for the first 10^6^ time steps. Nucleation indicates that the system is initialised in a metastable state, namely at a point of the phase diagram between the spinodal and the binodal; absence of nucleation indicates instead that the system is unstable and the relaxation process is a standard spinodal decomposition^30,46^. In the latter case (*ε* > 7 *kT*) the average number of particles in a cluster scales approximately as ∼ *t*^0.6^, corresponding to a cluster diameter growing with exponent *α* ≃ 0.6/2 = 0.3. Provided that *n*_B_*/n*_A_ stays close to unity, this is true also for *q* = 3 and 5, as shown by Fig. 7, where values of the cluster diameter growth exponent *α* obtained from fitting cluster size curves at late times are reported. A power law behaviour is a sign of self-similarity, which typically characterises relaxation to equilibrium of quenched interacting system (here quenching means initialising the system in the vapour state, but running the simulation at temperature lower than critical, where the equilibrium state exhibits phase separation).

**FIG. 6.**
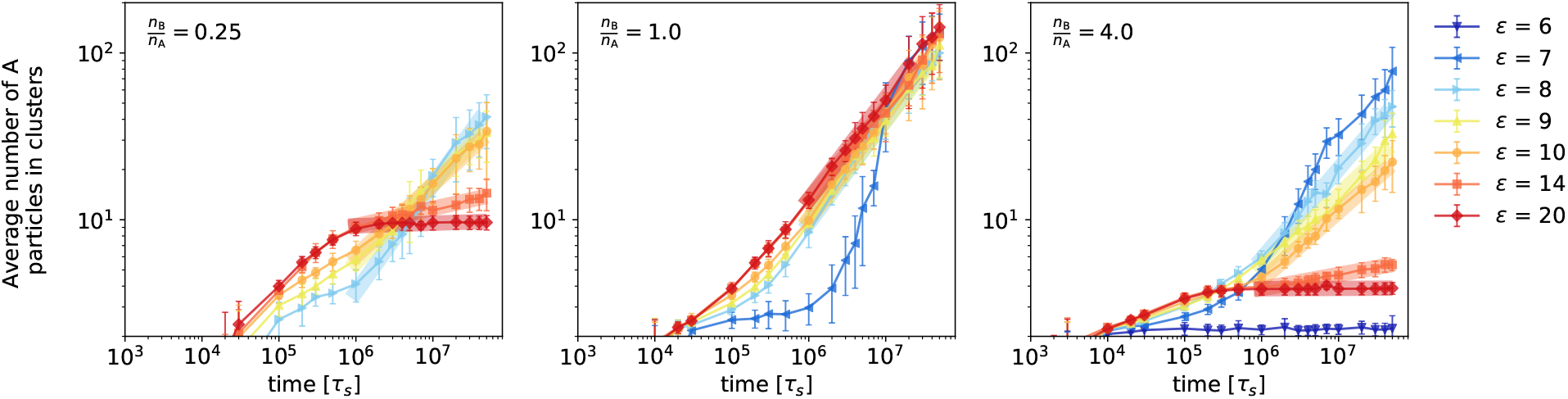
Time evolution of the average cluster size, as plotted in Fig. 3 and as defined in the text and in Methods, for *q* = 4. From left to right, the molar ratio *n*_B_*/n*_A_ is 1/4, 1 and 4. Bond energies *ε* are in units of *kT*. Thick lines show the intervals of linear behaviour from which we extract growth exponents for Fig. 7. Energy not shown do not give clusters.

**FIG. 7.**
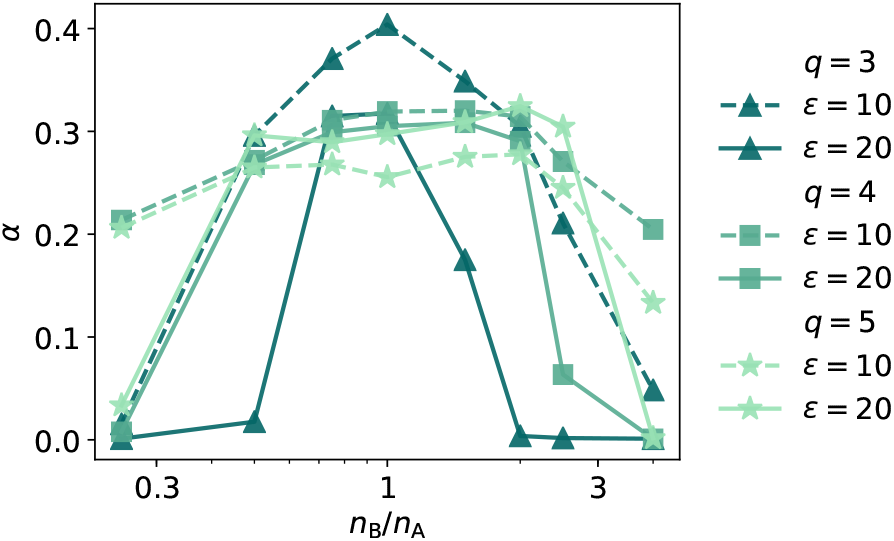
Growth exponent *α* measured from linear fit of cluster growth curves in time (Fig. 6). At concentration ratio close to 1, *α* ≃ 0.3 for most curves. Away from this regime, growth is hindered: coarsening is slowed down for *ε* = 10 *kT* (dashed lines) and completely halted for *ε* = 20 *kT* (continuous lines).

For low or high concentration ratios, the aggregation kinetics is slower. At small *ε*, this might be due to larger nucleation times, but linear behaviour in Fig. 6 rather points at a slow down in the power-law growth rate, at least for the probed times. This is consistent with the fact that at large *ε*, coarsening halts completely after a given time. This observation reflects the more pronounced non-monotonicity observed at final time for large bond strength in Fig. 3. Fig. 7 captures the decay of exponent *α* for different valences, as the concentration ratio deviates from unity.

We highlight that the process we see is mostly due to merging of existing clusters and not due to clusters growing by monomer addition^29^ (i.e. condensation of free particles): the latter process occurs very rapidly, usually in the very first steps of the simulation. This is confirmed by the observation that the fraction of clustered A reaches steady state on a time scale smaller than those represented in Fig. 6 (*<* 10^5^*τ*_*s*_).

A value of the exponent *α* ≃ 0.3 calls for a comparison with values reported in the literature. For 2D solid binary mixtures, an exponent *α* = 1/5 was proposed when growth is dominated by cluster diffusion and coalescence, at temperatures well below the critical one^31,32^. The so-called Lifshitz-Slyozov mechanism, akin to Ostwald ripening and due to evaporation and recondensation of single particles, should instead give *α* = 1/3, irrespective of dimensionality^30,33^. A richer phenomenology has been reported for fluid systems, for which (still in 2D) a crossover from *α* = 1/3, to 1/2, to 2/3 is expected^30,34^, although values of *α* between 1/4 and 1/3 have also been observed when hydrodynamics is suppressed^49,50^. Our system is neither fully solid, because clusters can diffuse and coalesce, nor fully fluid, as hydrodynamic drag effects are negligible at our low densities. In addition the morphology of the clusters and the stoichiometry can affect growth^30,47^.

We provide a simple theoretical argument, analogous to the one proposed in Ref. 51, showing why *α* in our case should be close to 1/4. We assume that the system exhibits dynamic scaling (i.e. self-similarity), that the probability of coalescence upon collision does not depend on time (which is plausible when growth is not arrested), that all clusters have equal size *M* at a given time, for simplicity, and that the cluster density *ρ*_*M*_ (number of clusters per unit surface at a given time) is small compared to the density of molecules within clusters. Now, since the total number of molecules is conserved, *ρ*_*M*_ = *ρ*_1_*/M* where *ρ*_1_ is the total number of molecules per unit surface. The average distance between two clusters at a given time is then 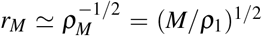. Given that the diffusion coefficient scales linearly with the mass, i.e. *D*_*M*_ = *D*_1_*/M*, the typical time *t*_*M*_ it takes for two clusters of size *M* to collide under the only effect of diffusion can be estimated as 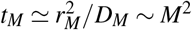. Neglecting fractal effects, the mass of the clusters scales as the square of its linear dimension *ℓ*_*M*_ (its diameter, for instance): 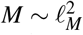, hence 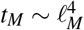. Since the system is self-similar, this relation must hold for any length in the system and at any time, giving *α* = 1/4.

### D. Size control is retained in fluid clusters

The clusters of the simple model we analysed so far are essentially solid-like: especially for *q* = 3 and 4, they are crystalline and their internal conformation can only change by evaporation of material and condensation of new material at a different place. This means defects can be eliminated only if they occur at the cluster boundaries. We show that the size-controlling mechanism we propose can be physiologically relevant, by making our clusters fluid. Biological condensates are indeed liquid-like even in 2D – although the availability of cross-linking molecules has been shown to reduce their internal diffusivity^17^.

To obtain fluidity, we add a generic isotropic attraction between any two particles (homo- and heterotypic), parametrised by its strength *ε*_*a*_ and range *R*_*a*_ (see Methods, Eq. (3)). Such attraction allows for local rearrangement once A-B bonds break: in this case particles can move around without having to leave the cluster, as some cohesion energy is still provided by isotropic interactions. The diffusivity of particles within clusters (internal diffusivity *D*_int_) is a measure of cluster fluidity. This is estimated, at late times, in the top plot of Fig. 8, as a function of bond strength *ε* and isotropic attraction *ε*_*a*_, for a chosen range *R*_*a*_. While in the large *ε* regime particles are locked by unbreakable bonds, there is a window where diffusivity is large. This occurs at small enough *ε*, so that bonds can be broken, yet large enough *ε*_*a*_ so as to guarantee cluster formation and stability.

**FIG. 8.**
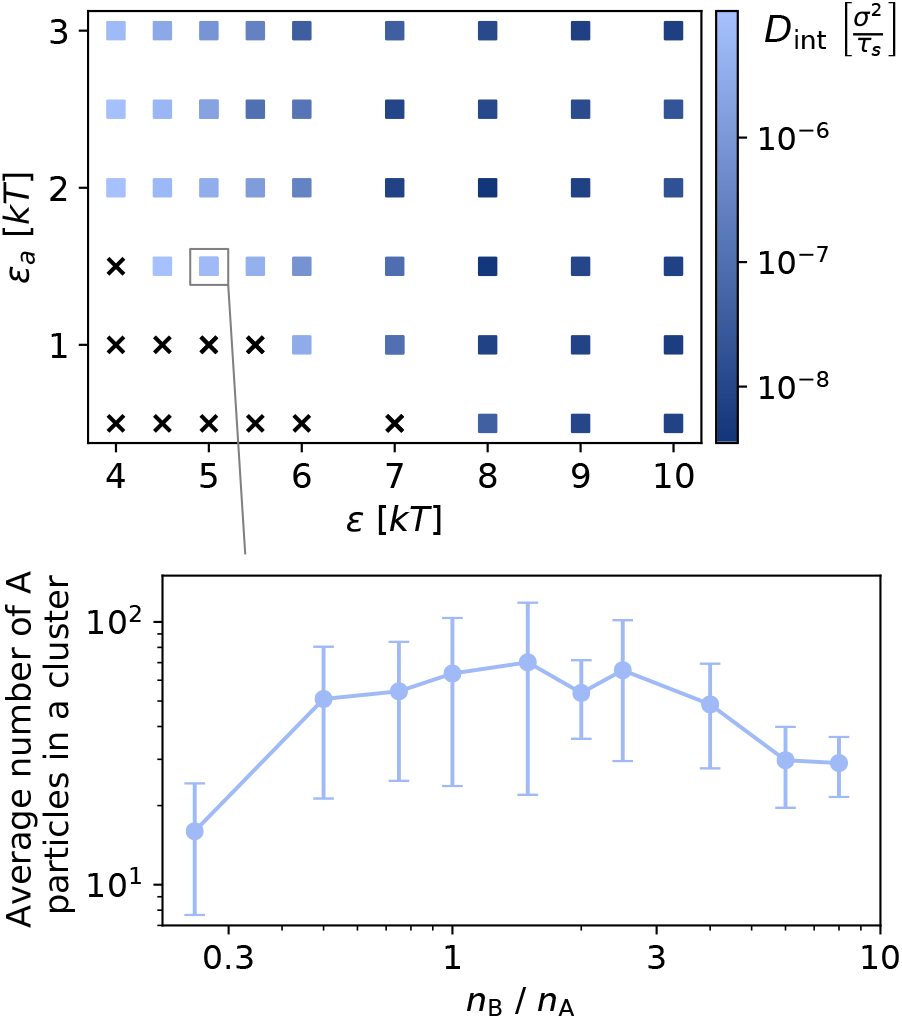
Top panel: the estimated internal diffusivity *D*_int_, for *q* = 5 and *R*_*a*_ = 2.0 *σ*, measured at *n*_B_*/n*_A_ = 1. Black crosses indicate that clusters do not form or are too volatile. *D*_int_ is computed by fitting the mean square displacement of particle pairs belonging to the same cluster in the time range 4 − 4.1 ×10^7^ *τ*_*s*_; pairs with one or both particles evaporating from the cluster within this time are discarded from the calculation. Bottom panel: an example of non-monotonic cluster size, measured at the end of the simulation, for a set of parameters where clusters are very fluid.

If the fraction of cluster energy provided by bonds stays comparable to the one provided by isotropic attraction, meaning that clusters would fall apart without specific interactions, unbound particles are more likely to leave the cluster than bound ones. We then expect to retain a signature of the size selection mechanism we discussed above. This is shown in the bottom plot of Fig. 8, featuring non-monotonic behaviour of the average cluster size with concentration ratio, in a case with fluid clusters. When the proportion of energy coming from isotropic attraction is too high (large *ε*_*a*_), this behaviour is lost.

## IV. DISCUSSION

Using a model based on patchy colloids with simple geometries, we studied a 2D binary mixture where specific heterotypic interactions are dominant. When the concentrations of the two species are comparable, clusters form and can in principle grow indefinitely. On the contrary, when one species is overabundant cluster growth is hindered, resulting in smaller clusters (Fig. 3). This happens because particles from the minority species tend to have all their binding sites saturated by particles of the majority species. Such capping phenomenon results in cluster surfaces being coated by same-kind particles, which cannot bind, thus preventing cluster coarsening. This phenomenon has been experimentally observed in biological condensates^16,17^ that do not belong to the client-scaffold category.

We show that such size-control process, although it can lead to stationary states within observation time (Fig. 6), is not of obvious thermodynamic origin, as instead recently found for analogous processes in different systems^19,52^. On the contrary, it can be driven by a kinetic trap whose depth increases with bond strength and unevenness in concentrations. This is reflected by a reduction in growth exponents, from ≃ 0.3 to 0, upon increase of the latter two quantities (Fig. 7). While large concentration unevenness decreases the probability for two clusters to coalesce, through the coating mechanism discussed above, strong bonds prevent bond rearrangement, necessary to reach equilibrium and eliminate defects. However, we still observe significant concentration-dependent size control for any value of bond strength.

Yet another way to control cluster size is by modulating valence. This strategy allows cells to tune properties of physiological condensates^2,53^, particularly on membranes^54^, but is also the functioning mechanism of artificial optogenetic tools used to study phase-separation^7,55^. We see that increasing valence broadens the range of concentration ratios at which clusters form. Full capping can indeed only occur when *n*_B_*/n*_A_ ≥*q* (or ≤1*/q*), as confirmed by our analysis of cluster size and growth exponent (Figs. 3 and 7).

Similarly to cluster size, concentration ratios and bond strength also affect the internal connectivity of the cluster: the connectivity of one species is obviously promoted by increasing the concentration of the crosslinking species, until it saturates (Fig. 2). The connectivity of the majority species decays with its concentration though. Optimal cluster connectivity is reached at equal concentration ratios. No matter the obvious limitations of the model, where spurious geometric effects are present, these considerations are mirrored by diffusivity measurements in presence of heterotypic interactions only^17^.

In conclusion, we highlight how clustering curves as a function of concentration ratio might be a way to infer properties on the molecular scale. Our results suggest that if clustering is non-monotonic, specific heterotypic interactions are possibly responsible. The more pronounced the non-monotonicity, the stronger are specific bonds with respect to generic (isotropic) interactions. The latter, however, might be present if the cluster appears macroscopically fluid. Finally, the kinetics of cluster growth carries likewise relevant information: a decrease in growth exponent with concentration ratio might as well be a signature of dominantly heterotypic interactions.

## ACKNOWLEDGMENTS

We thank Longhui Zeng and Xiaolei Su (Yale University) for bringing the topic to our attention and for useful comments. This work has received funding by the European Research Council under the European Union’s Horizon 2020 research and innovation programme (ERC grant No. 802960 and Marie Skłodowska-Curie grant No. 101034413). We are grateful to the UK Materials and Molecular Modelling Hub for computational resources, which is partially funded by EPSRC (Grant No. EP/P020194/1 and EP/T022213/1). We acknowledge support from IST Austria and from the Royal Society (Grant No. UF160266).

## APPENDIX

### CALCULATION OF CONSTRAINED FREE ENERGY

The partition function of a macroscopic state where the maximum number of bonds are formed (*qn*_A_) and there is only one giant cluster of whatever shape is the following:

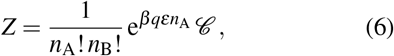

where *β* = (*kT*)^−1^ and 𝒞 is the number of different microscopic configurations corresponding to the same macroscopic state we are constraining on.

We group configurations based on the number *n*_Bc_ of B particles belonging to the cluster; the free B particles are then *n*_Bf_ = *n*_B_ − *n*_Bc_. We have

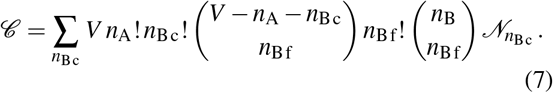

The seven factors of this sum represent, respectively: positions of the cluster in a lattice with *V* sites, permutations of the *n*_A_ A particles in the cluster, permutations of the *n*_Bc_ B particles in the cluster, positions of the *n*_Bf_ free B particles (to be chosen among the sites not occupied by the cluster), permutations of the *n*_Bf_ free B particles, ways to chose which B particles are free and which belong to the cluster, and finally 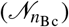 number of different cluster shapes (including rotations) that embed *n*_Bc_ particles in the cluster.

The B particles in the cluster can have either more than one bond (we say that they belong to the bulk) or only one bond (they form the boundary). The bulk contains always one B particle per A particle (see top-right sketch in Fig. 5), so there are *n*_A_ B particles in it. The number of boundary particles, instead, is proportional to the perimeter of the cluster bulk. In summary, the number of B particles in the cluster is *n*_Bc_ = *n*_A_ + *p*, where *p* is the semiperimeter of the shape formed by the 2*n*_A_ particles in the bulk. Eq. (7) is then interpretable as a sum over all possible perimeters, and 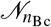 as the number of different shapes of semiperimeter *p* = *n*_Bc_ −*n*_A_ that can be built clustering 2*n*_A_ particles.

The statistics of connected shapes on lattices, so-called lattice animals or polyominos, has been a long studied subject^45^. 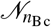 can be written as the product of the total number *a*_*s*_ of shapes of size *s* = 2*n*_A_ (scaling as 4.0^*s*^*/s*)^44,45^ and the perimeter distribution *f*_*p*_ (i.e. the fraction of shapes of size *s* that have semiperimeter *p*). To our knowledge a rigorous estimate of the asymptotic behaviour of the latter quantity does not exist, so we extracted *f*_*p*_ for *s* up to 48 from exact tabulated enumerations of lattice animals^42,43,56^. Fig. 9 shows that semiperimeters, rescaled by 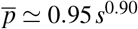, seem to follow approximately a normal distribution with mean *μ* = 1 and variance *σ* = 0.06.

**FIG. 9.**
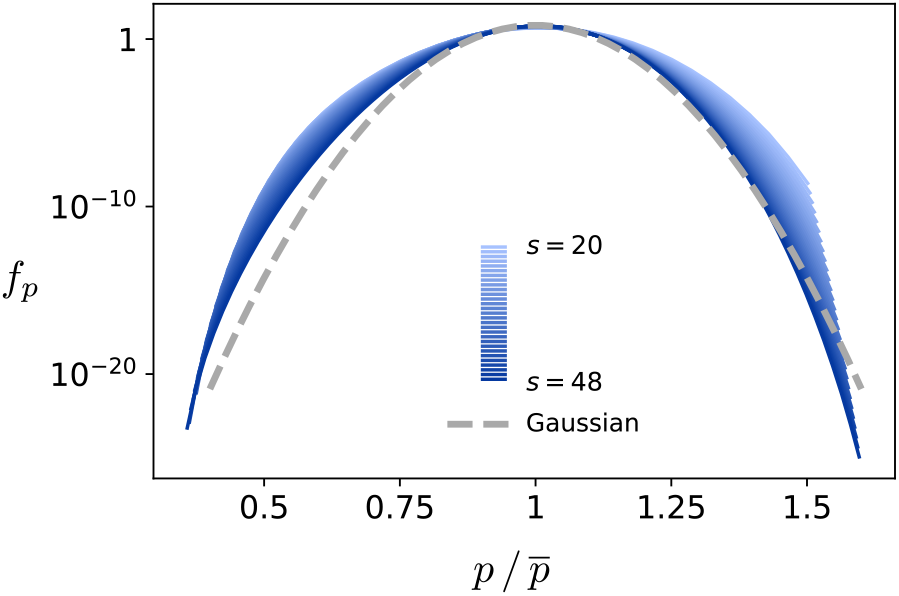
Distribution of semiperimeters, as a function of rescaled semiperimeter *p*. The rescaling factor 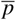 is defined in the text. The curves for sizes *s* from 20 to 48 (obtained from Ref. 56) approximately collapse on a single curve that resembles a Gaussian. Note that the vertical scale is logarithmic.

In summary, Eq. (7) can be rewritten as

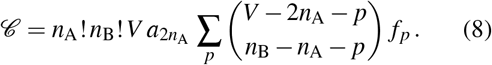

In the sum, notice the competition between the second term, representing “shape” entropy (more shapes are available at larger perimeter), and the first term, representing translational entropy of free B particles (larger cluster perimeter means fewer B particles in solution). Using Stirling’s approximation and passing to the continuous limit for *f*_*p*_, we obtain

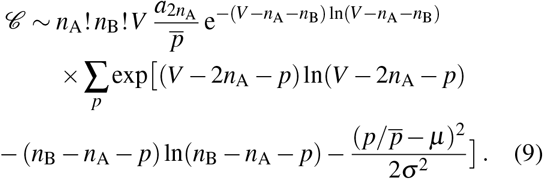

We verified numerically that the Gaussian term only plays a role at very high densities, not relevant in our work. The dominant term of the sum is then the smallest perimeter one. To leading order, this amounts to setting 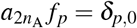 (Kronecker delta) in Eq. (8). By doing so, one recovers Eq. (4), which refers to a circular cluster.

